# First report on chromosome number and genome size of rare endemic birch *Betula klokovii* Zaverucha from Ukraine

**DOI:** 10.64898/2026.10.14.759152

**Authors:** Andrii Tarieiev, Grit Winterfeld, Laura Freisleben, Andrii Shtogun, Martin Röser

**Affiliations:** Institute of Biology, Geobotany and Botanical Garden, Martin Luther University Halle-Wittenberg, Am Kirchtor 1, 06108 Halle (Saale), Germany; National Nature Park “Kremenets Mountains”, Osovytsia Str. 12, 47003 Kremenets, Ukraine

**Keywords:** karyology, chromosome number, genome size, flow cytometry (FCM), *Betula*, endemic species, Ukraine

## Abstract

This study presents the first report on chromosome number and genome size estimate for rare endemic birch species *Betula klokovii* Zaverucha from Ukraine. Karyological analysis of metaphases revealed a total of 56 chromosomes, confirming a tetraploid cytotype (2*n* = 4*x* = 56). Concurrently, flow cytometry (FCM) was utilized to conduct a precise genome estimation, yielding a nuclear DNA content of 2C = 1.93–1.99 pg for fresh material and 2.15–2.53 pg for herbarium. In addition, we identified one seedling with lower DNA content of 2C = 1.46 pg that most likely corresponds to F_1_ triploid hybrid between *B. klokovii* and *B. pendula* Roth. The results contribute significantly to our understanding of genomic evolution within the genus *Betula* L. and provide important insights into the taxonomy and conservation genetics of rare European birch taxa.

## Introduction

Birches are common trees and shrubs in the Northern Hemisphere, where the genus *Betula* L. comprises approximately 65 species (Ashburner & McAllister, 2016), with estimates ranging from 30 (de Jong, 1993) to 89 (Govaerts, 2024), worldwide. The genus is highly diverse and comprises many polyploids and hybrids, and hybridization also occurs between species with different ploidy levels (Johnsson, 1945; Anamthawat-Jónsson, 2012; Ashburner & McAllister, 2016; Wang, 2016). The basic chromosome number in birch trees is 14, the range extends from 2*n* = 2*x* = 28 to 2*n* = 12*x* = 168 (Rice & al., 2015; Ashburner & McAllister, 2016). The genome size (1C) varies from 0.44 pg for diploid *B. nigra* L. to 2.56 pg for dodecaploid *B. megrelica* Sosn. and 2.59 pg for *B. fargesii* Franch. without a known ploidy level (Wang & al., 2016; Leitch & al., 2019). The ploidy levels in birch correlate with the climatic niche breadth: tetraploid birch taxa exhibit the largest geographic range size and the widest climatic niche breadth, with *B. pubescens* Ehrh. displaying the largest area of occupancy while high polyploids usually have a narrow distribution (Yan & al., 2025).

In birch taxa, nuclear DNA content directly correlates with ploidy levels, enabling indirect ploidy estimation. However, many birch taxa are still lacking both cytometric and karyological data.

*Betula klokovii* Zaverucha is a rare, endemic birch species found only on two mountains – Strakhova and Maslyatyn – near Kremenets in Ternopil region of Ukraine. Klokov’s birch was initially described by Borys Zaverucha (Zaverucha, 1964) as a 5–10 m tall tree, with 1–2 stems and white bark that does not peel off and has a specific dotted pattern on old trunks (Fig. 1). Young twigs are non-pendulous, with cherry-brown bark color, densely or sparsely pubescent. Buds are sticky, pointed, 4–6 mm long. Leaves are rhombic-ovate, or oblong-ovate with a cuneate base, 25–55 mm long, and 15–40 mm wide. The adaxial surface is dark green, glabrous, whereas the abaxial surface is pale greyish green, somewhat pubescent (Zaverucha, 1964). The type specimens of *B. klokovii* including holotype KW000006416 as well as isotypes KW000006417–KW000006421, and original material (specimina authentica) KW000006422–KW000006425 are stored in the National Herbarium of Ukraine (KW) (Olshanskyi & al., 2016).

**Fig. 1.**
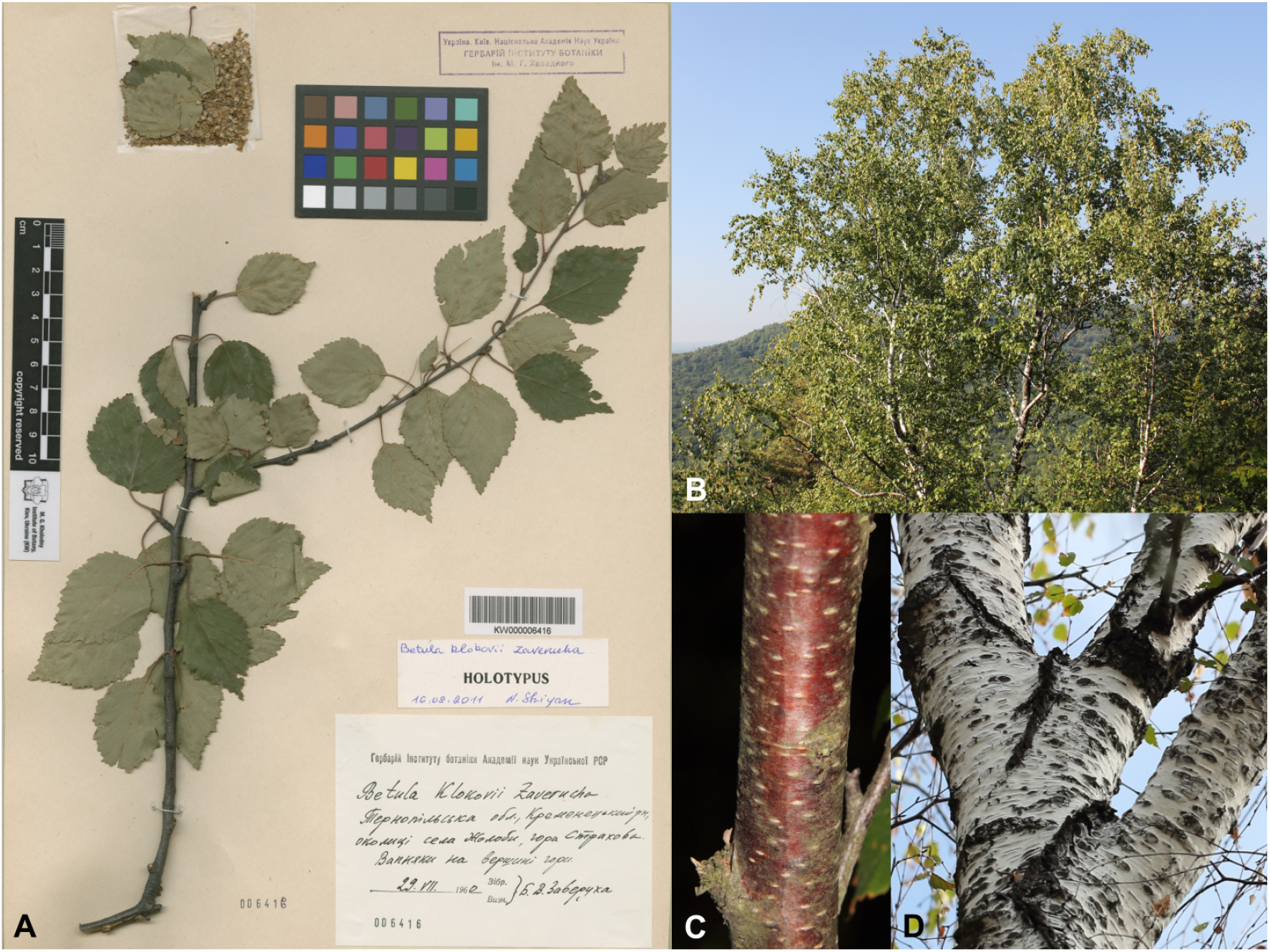
*B. klokovii* Zaverucha: A – holotype stored in the National Herbarium of Ukraine (KW000006416); B – general habitus of Klokov’s birch; C – bark on the young twig; D – stem of the tree with characteristic dotted pattern

The very small natural habitat and very small number of mature trees pose a significant threat to this species. Consequently, it is protected at both local and state levels in Ukraine (Kagalo & al., 2009), and is listed as Critically Endangered on the IUCN red list (Rivers & Tarieiev, 2015). Both populations are located on the territory of Natural Nature Park (NNP) “Kremenets Mountains”. In order to facilitate conservation efforts, Mr. Tarieiev together with the staff of NNP “Kremenets Mountains” performed morphological identification of the trees and collected GPS coordinates (in 2014–2019). Finally, a special nursery for planting young trees from seeds was established in 2017. The distribution of the trees and the location of the nursery are shown on Fig. 2.

**Fig. 2.**
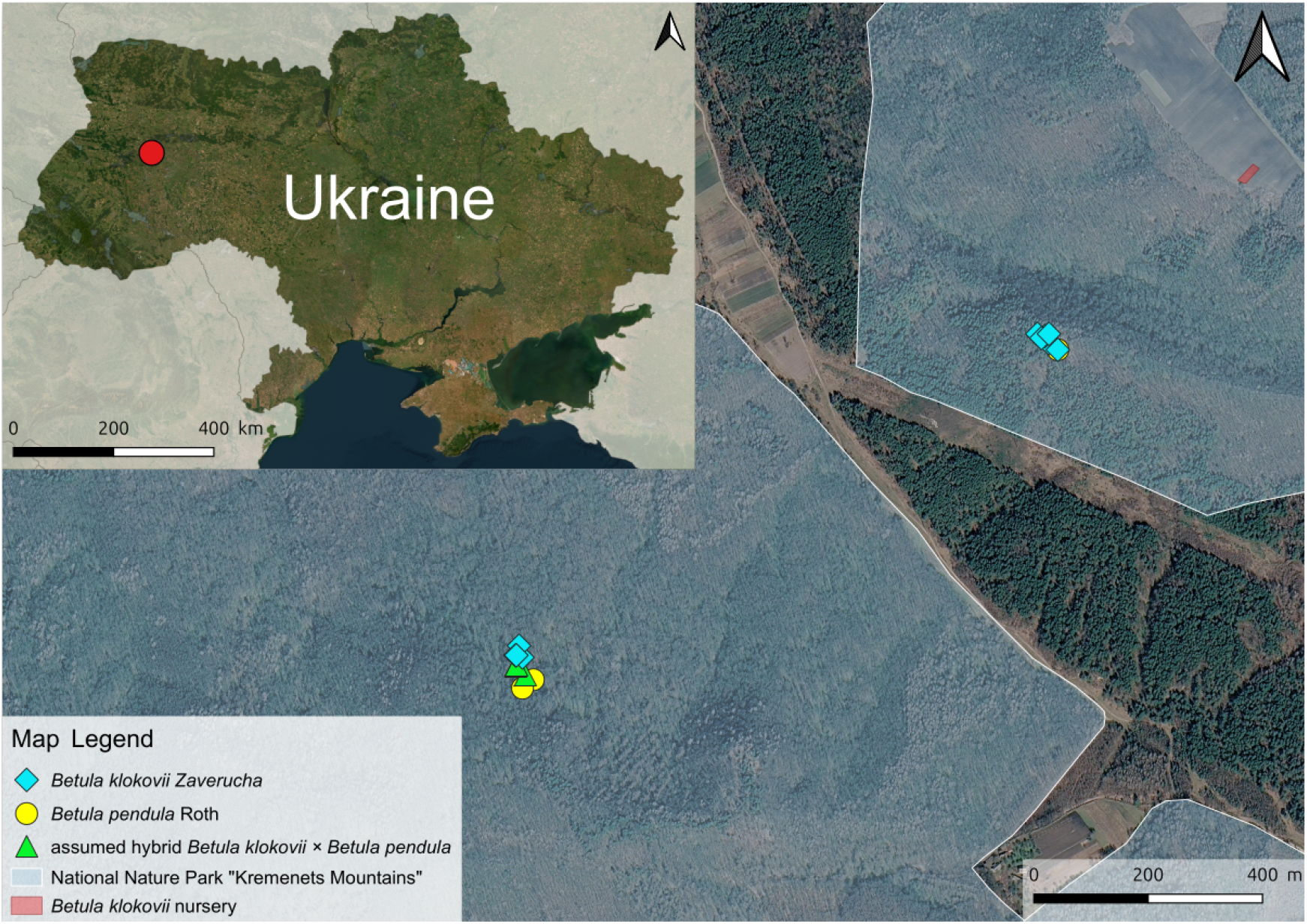
Distribution map of *Betula klokovii* Zaverucha. Created in QGIS 3.44.15-Solothurn using ESRI and Google Satellite Map

Although *B. klokovii* has been hypothesized to share a tetraploid cytotype due to its close phylogenetic relationship with *B. pubescens* (Zaverucha, 1964; Walters, 1993; Tzvelev, 2002; Tarieiev & al., 2025), the precise data confirming its chromosome number and genome size are still lacking. Hence, we conducted both karyological and flow cytometric (FCM) analyses on herbarium material and seedlings grown from seeds collected from mature trees that morphologically correspond to the protologue (initial description) of *B. klokovii*.

## Material and methods

### Plant material

Study was conducted on seedlings grown from seeds collected by Mr. Andrii Shtogun in 2025 from multiple mature trees within the National Nature Park “Kremenets Mountains” that morphologically correspond to the original description of the species.

In addition, we used silica gel dried herbarium material collected in 2019 from trees corresponding to the protologue of *B. klokovii* (herbarium specimens are deposited in GOET, detailed information is provided in the Supplementary Table 1).

After cold stratification, the seeds were germinated on DKW medium (Duchefa Biochenie, Netherlands) solidified with 0.8% agar (Carl Roth, Germany). Obtained seedlings were subsequently harvested and utilized for both karyological and FCM analysis.

### Karyological analysis

The seedlings were washed with tap water a few times, treated in 0.002 M water solution of 8-hydroxyquinolin (Merck, Darmstadt, Germany) at RT for 3h to accumulate metaphases, fixed absolute ethanol/ glacial acetic acid (AppliChem, Darmstadt, Germany) 3:1 v/v for 3h at RT and stored in absolute ethanol at -20°C until preparation. The seedlings were macerated (Schwarzacher & al., 1980) in enzymatic mixture of cellulase from *Aspergillus niger* (liophylized powder, 4310u/g, Calbiochem, La Jolla, CA) and pectinase P-5146 from *Aspergillus niger* (solution in 40% glycerol, Sigma, USA) (1%:10%, w/v) at 37°C for 30 min. The seedlings were squashed on slides in a drop of 45% propionic acid (Carl Roth, Karlsruhe, Germany) with 2% carmine (Merck, Darmstadt, Germany) and covered with a cover slip. Photographs of metaphase chromosomes were taken on a Zeiss Axiophot microscope using a computer-assisted cooled Zeiss Axiocam 807 camera.

For karyotype analysis, a single metaphase plate displaying well-defined, sharp chromosomal contours was selected. Chromosomes were manually arranged and measured using CorelDRAW^®^ Graphics Suite X8. Digital measurements of chromosome length and constriction positions were processed in Microsoft Excel 2010 for subsequent calculations. The mean chromosome length was calculated as *x*CL = (L+S) / cn [µm], where L is the total length of the long arms, S is the total length of short the arms in a chromosome set, and cn represents the chromosome number. Chromosomes were numbered in descending order based on their total length. Structural classification was determined by centromere position according to Levan et al. (Levan & al., 2009), identifying chromosomes as either metacentric (m), submetacentric (sm), or subtelocentric (st). Putative homologous or homoeologous chromosomes were then paired, based on similarities in length and centromeric indices, numbered with Roman numerals and grouped according to a possible distribution as two diploid genomes.

### Genome size measurement

Absolute genome sizes (nuclear 2C values = holoploid genome sizes) were estimated on both fresh (seedlings) and herbarium material by propidium iodide flow cytometry (FCM) on CyFlow Ploidy Analyzer (Partec, Germany) with a Nd-YAG green solid-state laser (532 nm) using CyStain PI OxProtect reagent kit for nuclei extraction and DNA staining of nuclear DNA from plant tissues to determine absolute and relative genome size and ploidy level (Viruel & al., 2019; Boutte & al., 2020) (https://products.sysmex-partec.com/cystain/cystain-ploidy-dna-analysis/2964/cystain-pi-oxprotect#) according manufacturer’s instructions. The protocol is described in detail in (Winterfeld & al., 2025). All the measurements on herbarium material were performed three times on the same sample as suggested by (Loureiro & al., 2023). Seedlings were analyzed individually, but were pooled in pairs if the available plant tissue volume was insufficient for separate analysis. *Raphanus sativus* L. ‘Saxa’ with DNA content 2C = 1.11 pg, and *Glycine max* Merr. ‘Polanka’ with DNA content 2C = 2.50 pg (Doležel & al., 2007) were used as internal standards. Data evaluation was performed using the FCS Express v5 Flow (De Novo Software, USA). Within this study a total of seven measurements were conducted on seed samples, alongside five measurements on herbarium specimens.

## Results

### Chromosome count and karyotype analysis

Karyological examination of seven metaphase plates consistently revealed a chromosome number around 2*n* = 56. In particular, a chromosome count of 2*n* = 56 was confirmed in four cells, while the remaining three plates exhibited a slight variation ranging from 2*n* = 54 to 56. A representative metaphase plate showing 2*n* = 56 chromosomes, along with the corresponding idiograms arranged as two diploid (sub-)genomes A and A’, is presented in Figure 3. The calculated mean chromosome length was *x*CL = 1.04 µm.

**Fig. 3.**
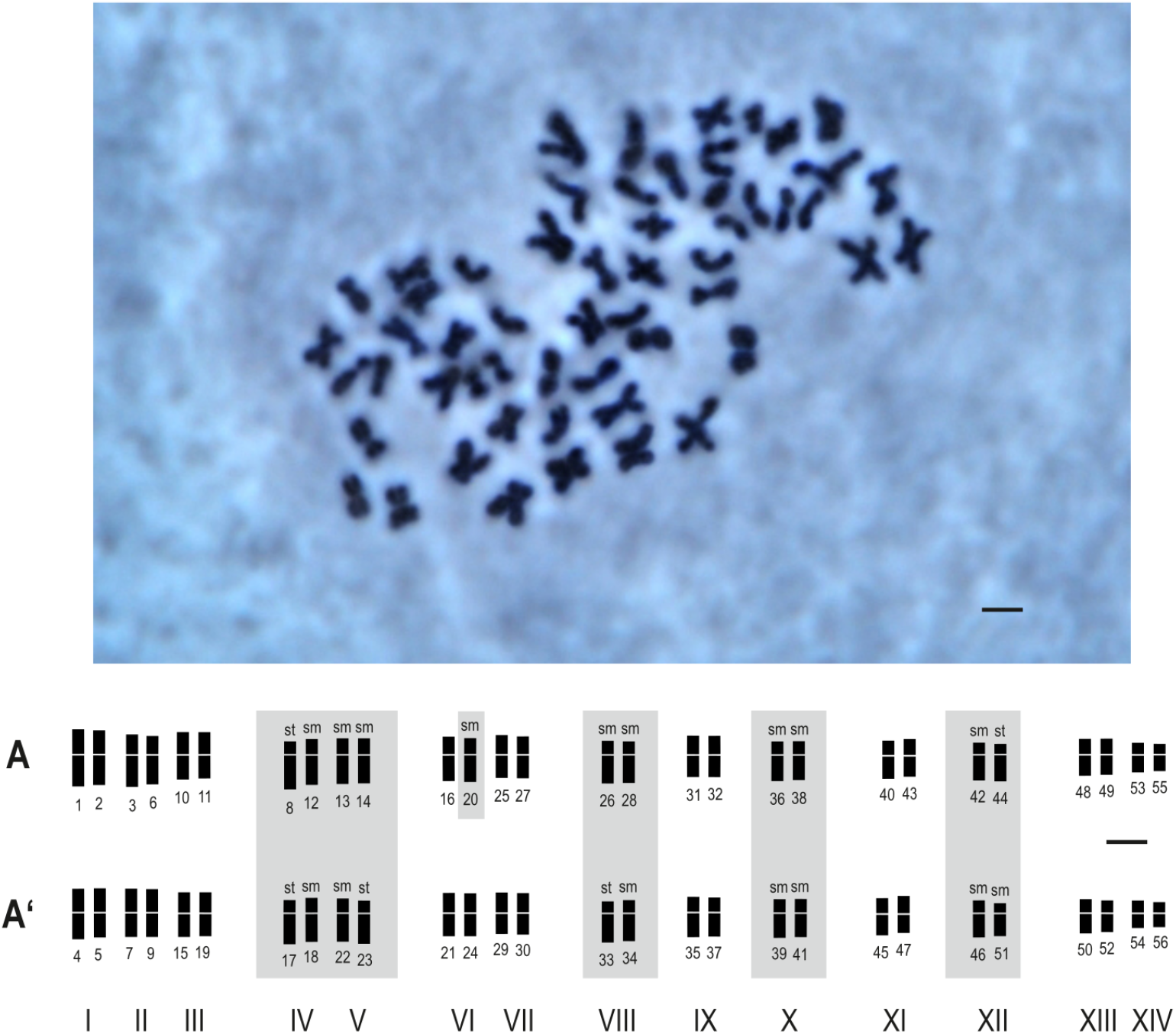
Somatic metaphase plate (top) and chromosome idiogram (bottom) of *Betula klokovii* with 2*n* = 4*x* = 56. Arabic numerals designate chromosomes ordered by individual length. Roman numerals indicate chromosome pairs of putative homologous or homoeologous groups. A and A’ represent the putative distribution into two distinct genomes. Chromosome morphology: sm, submetacentric; st, subtelocentric (both highlighted in grey); unmarked chromosomes are metacentric Scale bar = 1 µm.

### Genome size

Flow cytometric measurements yielded an absolute nuclear DNA content of 2C = 1.93–1.99 pg for fresh material (seedlings) and 2C = 2.17–2.26 pg for herbarium material. Between analyzed seedlings we identified one individual with a lower DNA content of 2C = 1.46 pg that most likely is a triploid hybrid between *B. klokovii* and *B. pendula* Roth. The details on measurements are provided in Supplementary Tables 1 and 2.

## Discussion

Both chromosome count and genome size estimations show that *Betula klokovii* is a tetraploid species. This result confirms earlier assumptions regarding its ploidy level (Zaverucha, 1964; Walters, 1993; Tzvelev, 2002; Tarieiev & al., 2025) which is frequently discussed alongside its close phylogenetic relationship to *B. pubescens*. Furthermore, our study provides robust evidence for hybridization between *B. klokovii* and *B. pendula*, which aligns with our earlier findings inferred from morphology (Tarieiev & al., 2025), and described cases of hybridization between di- and tetraploid birch species resulting in triploid hybrids. In particular, such a hybrid between *B. pendula* and *B. pubescens* was described as *B. ×aurata* Borkh. from Grübesberg in Thuringia (Borkhausen, 1800), and there are cytological studies on Icelandic triploid hybrids between *B. pubescens* and *B. nana* L. (Anamthawat-Jónsson & Thór Thórsson, 2003; Anamthawat-Jónsson & al., 2021). However, further investigations further investigations are necessary to determine the evolutionary role, frequency and the underlying genetic mechanisms of hybridisation between *B. klokovii* and *B. pendula*.

Although the usage of dried plant material for FCM is well-established (Suda & Trávníček, 2006a, 2006b), it works not for all plant taxa, and even rapid drying introduces additional error (Bainard & al., 2011).

All FCM genome size estimates of herbarium material showed higher DNA content than fresh samples. The same effect was already observed in *Fagus grandifolia* Ehrh., *Taraxacum officinale* F.H. Wigg., *Typha latifolia* L., *Urtica dioica* L., and *Leonurus cardiaca* L. (Bainard & al., 2011). The discrepancy in genome sizes between dried herbarium specimens and fresh material, along with the significantly higher dispersion observed in the dried material, could potentially be explained by cellular and DNA degradation during drying, and subsequent storage. Other factors like number and amount of secondary metabolites can also contribute (Loureiro & al., 2023). To address this discrepancy, flow cytometric (FCM) analyses should ideally be performed on fresh material collected from adult trees. Unfortunately, such material was unavailable for the present study.

It is important to note that experiments were conducted on seedlings that cannot be reliably identified morphologically, as the protologue describes only adult trees. Given that birches are well-known for frequent interspecific hybridization (Johnsson, 1945; Ashburner & McAllister, 2016; Wang & al., 2016; Bona & al., 2018; Anamthawat-Jónsson, 2019), the presence of hybrid individuals among the seedlings cannot be entirely ruled out. Notably, our FCM measurements revealed that one seedling possessed a genome size of 2C = 1.46 pg. This individual is most likely a triploid F_1_ hybrid resulting from a cross between the tetraploid *B. klokovii* and diploid *B. pendula*. Since reliable morphological identification of birches in juvenile state is very challenging to impossible, FCM gemone size extimates can be suggested as a cost efficient method for screening trees and saplings to identify F_1_ hybrids between Klokov and silver birches. Such information will not only help to understand interspecific hybridization and introgression better but also relevant for developing conservation measures.

To gain deeper insights into the evolutionary dynamics of *B. klokovii*, separate and comprehensive studies are required. Future research should integrate karyological analyses, FCM genome size estimations, fluorescence/ genomic in situ hybridization (FISH/GISH), and whole-genome sequencing across a larger number of individuals, ideally encompassing the entire population.

## Conclusion

Both chromosome counts and flow cytometric (FCM) genome size estimations confirm that Ukrainian the endemic species *Betula klokovii* is tetraploid. Somatic metaphase plates obtained from the seedlings of *B. klokovii* consistantly reveal a chromosome number of 2*n* = 4*x* = 56. The chromosomes were morphologically classified as either metacentric (m), submetacentric (sm), or subtelocentric (st). Genome size estimates yielding a value of 2C = 1.93–1.99 pg for seedlings, and 2C = 2.17–2.26 pg for herbarium material. Notably, one seedling exhibited a significantly lower DNA content (2C = 1.46 pg) and most likely represents a triploid F_1_ hybrid between the tetraploid *B. klokovii* and the diploid *B. pendula*.

## Author contribution

AT – initial idea of the study, partial labwork, planting seeds, writing initial draft, editing the manuscript; GW – cytogenetics and karyology, editing the manuscript; LF – laboratory assistance, flow cytometry measurements, editing the manuscript; AS – seed collection, editing the manuscript; MR – funding acquisition, editing the manuscript.

