## Supplementary Table 1 for "First report on chromosome number and genome size of rare endemic birch *Betula klokovii* Zaverucha from Ukraine"

**Supplementary Table 1.** Herbarium specimens used for FCM

| Taxon | Herbarium sample identifier | Sample identifier | Collection place | GPS | Collection date | Collectors | Reference | Genome size (x̄), pg (FCM) | SD | SE |
|---|---|---|---|---|---|---|---|---|---|---|
| *B. klokovii* | GOET072790 | UA-KM-6Bk | mt. Maslyatyn, NNP “Kremenets Mountains”, near Kremenets, Ternopil region, Ukraine | N50°04.791' E025°39.690' | 16.07.2019 | A. Tarieiev, A. Shtogun, I. Dovhaniuk | *Raphanus sativus* L. ‘Saxa’ | 2.19 | 0.0495 | 0.035 |
| *B. klokovii* | GOET072788 | UA-KM-8Bk | mt. Maslyatyn, NNP “Kremenets Mountains”, near Kremenets, Ternopil region, Ukraine | N50°04.793' E025°39.680' | 16.07.2019 | A. Tarieiev, A. Shtogun, I. Dovhaniuk | *Raphanus sativus* L. ‘Saxa’ | 2.22 | 0.0208 | 0.012 |
| *B. klokovii* | GOET072793 | UA-KS-7Bk | mt. Strakhova (on the slope near the top), NNP “Kremenets Mountains”, near Kremenets, Ternopil region, Ukraine | N50°05.087' E025°40.462' | 15.07.2019 | A. Tarieiev, A. Shtogun, I. Dovhaniuk | *Glycine max* Merr. ‘Polanka’ | 2.26 | 0.0212 | 0.015 |
| *B. klokovii* | GOET072791 | UA-KS-10Bk | mt. Strakhova (on the top), NNP “Kremenets Mountains”, near Kremenets, Ternopil region, Ukraine | N50°05.096' E025°40.445' | 15.07.2019 | A. Tarieiev, A. Shtogun, I. Dovhaniuk | *Raphanus sativus* L. ‘Saxa’ | 2.17 | 0.0067 | 0.0115 |
| *B. klokovii* | GOET072785 | UA-KS-11Bk | mt. Strakhova (on the top), NNP “Kremenets Mountains”, near Kremenets, Ternopil region, Ukraine | N50°05.092' E025°40.451' | 15.07.2019 | A. Tarieiev, A. Shtogun, I. Dovhaniuk | *Raphanus sativus* L. ‘Saxa’ | 2.26 | 0.0929 | 0.0536 |
