## Supplementary Table 2 for "First report on chromosome number and genome size of rare endemic birch *Betula klokovii* Zaverucha from Ukraine"

**Supplementary Table 2.** FCM measurements of seedlings

| Seedling | Taxon | Reference | Genome size, pg (FCM) | x̄ | SD | SE |
|---|---|---|---|---|---|---|
| 1 | *B. klokovii* | *Glycine max* Merr. 'Polanka' | 1.93 | 1.96 | 0.0112 | 0.0273 |
| 2 | *B. klokovii* | | 1.95 | | | |
| 3 | *B. klokovii* | | 1.93 | | | |
| 4 | *B. klokovii* | | 1.95 | | | |
| 5 | *B. klokovii* | | 1.99 | | | |
| 6 | *B. klokovii* | | 1.99 | | | |
| 7 | *B. klokovii* × *B. pendula* | | 1.45 | – | | |
